# Novel MRI-based Biotypes of Human Neurodevelopment associated with Impulsivity performance

**DOI:** 10.1101/2024.11.11.623131

**Authors:** Zicong Zhou, Miguel A. Rivas-Fernandez, Shana Adise, Luis Colon-Perez

## Abstract

**Introduction:** Adequate neurodevelopment is critical to the proper advancement of cognitive and emotional processing; however, variation in neurodevelopment has emerged as a potential risk factor for subsequent drug use. The Adolescent Brain Cognitive Development (ABCD) Study is an ongoing and longitudinal study to determine the developmental trajectories of the brain and how experiences impact brain development, which makes it an ideal framework study to determine proper neurodevelopmental trajectories associated with healthy executive function behavioral development and their neurobiology.

**Method:** Based on the availability of ABCD (Release 5.1) tabulated data, we selected 1368 subjects with neuroimaging records on multiple time points. To focus on the frontal cortex, we preprocess 20 ROIs (including ten regions with right and left cerebral hemispheres) cortical thickness measurements for SuStaIn. Once SuStaIn determined the potential control and phenotype (subtypes) group, we conducted t-tests between the control and phenotype groups on their delay discounting measurements. To overserve possible trajectories, we also investigated the correlation of the thickness measurements with delay discounting under the suggested phenotype by SuStaIn.

**Results:** SuStaIn identified four subtypes and one potential control group, validated by cross-validation test as the optimal solution for balancing model accuracy and complexity. Among the t-tests on delay discounting results between the control group and each subtype, subtype1 tends to show low significance on the 3-month indifference point (p=0.067), subtype 2 identifies a significant reduction in impulsivity on the 1-month indifference point (p=0.017), subtype 4 stands significant reduction at on 3-month indifference point (p=0.0092) and significance on 1-year indifference point (p=0.00418). For the correlations between each selected ROI, we identified a negative correlation (R=-0.18, p=0.0095) between the thickness of the left hemisphere caudal-anterior-cingulate and delay discounting indifference point at 3 months.

**Conclusion:** We successfully identified four subtypes of the ABCD, which may be putative abnormal trajectories of executive function accelerated development. Three of the four subtypes reflect impulsive behavioral phenotype, and one displays an association with cortical thinning. Overall, we present evidence that there are putative neurobiological signs of potential executive function disruption as early as nine years of age.

## Introduction

Adequate neurodevelopment is critical to the proper advancement of cognitive and emotional processing; however, variation in neurodevelopment has emerged as a potential risk factor for subsequent drug use. There is not much direct evidence of the brain’s regional foci and trajectories that predispose one to risky behavior and substance use; therefore, it is imperative to understand neurodevelopment and its associated risks for substance abuse during adolescence. Adolescence is a period in which the brain undergoes substantiative changes, and particular trajectories of these processes may increase the future risk for mental health issues like executive function deficits and altered cognitive control, which are risk factors for later substance use. Some examples of the active neural developmental processes during adolescence are changes in brain anatomy (cortical thickness and volumes),^1^ synaptic density,^2^ myelination,^3^ functional connectivity,^4^ and structural connectivity.^5^ The development of these trajectories is complex and involves many concurrent and independent stages modifying brain structure, and this process takes years to refine.^6^ During this process, there may be underlying differences in the development of neurocircuitry that may predispose some to become more impulsive and engage in substance use, but the complete profile of these neural signatures has not been well-identified, particularly those centered around frontal lobe structures.

The frontal lobe and its associated cognitive functions are essential for proper development.^7^ The frontal lobe is the epicenter of executive function, decision-making, and cognitive control, critical for healthy behavior and protection against risky behavior, especially those associated with substance use.^8–10^ Dysfunction in executive function and cognitive control leads to limitations in the capacity to organize goal-oriented behavior, leading drug users to seek and consume drugs despite the harmful consequences of their use.^11^ Dysfunction of the frontal cortex and its associated circuitry is associated with several neuropsychiatric disorders, including functions associated with the risk of drug use.^12^ Neuroanatomically, larger PFC volume and greater PFC thickness were associated with better executive performance.^12^ Also, high impulsivity is associated with a higher susceptibility to cocaine self-administration in rats.^13^ In fact, over 50 years of research and inquiry into executive function and cognitive processes are now associated with frontal lobe function, which is beyond the scope of this introduction. Nonetheless, it is accepted that executive function and cognitive control alterations are associated with early life adversity, stress, environmental and genetic factors. However, the neurobiological features during adolescence that predispose executive function and cognitive control alterations have not been elicited.

Machine learning (ML) techniques have emerged as an integral tool in research, from physics to neuroscience. ML allows researchers to utilize computers to identify complex patterns from datasets, enabling us to interpret data and determine relationships that otherwise would be difficult to identify. One model of interest for this application is the Subtype and Staging Inference (SuStaIn). This model is designed to determine biological subtypes with similar temporal trajectories (i.e., stages) from neuroimaging data. The biological subtypes (biotypes) represent an unbiased neural-derived phenotype. SuStaIn modeling has successfully identified two schizophrenia biotypes with structural MRI: the first starts with alterations in the hippocampus and the second with cortical (primarily, insula) alterations.^14^ In Alzheimer’s disease, SuStaIn uncovers three biotypes: one with foci in the hippocampus and amygdala, a second in the nucleus accumbens, insula and cingulate, and a third in the pallidum, putamen, nucleus accumbens, and caudate.^15^ While in multiple sclerosis, SuStaIn identifies three biotype groups: one subtype originates in the occipital and parietal cortex, a second starts in the white matter, specifically the cingulate bundle and corpus callosum, and a third on lesions through the brain.^16^ SuStaIn has been successfully deployed to identify potential regional nuance to disease development. However, it has rarely been followed up to determine the significance of such trajectories nor the predictive value of such biotypes in development and subsequent disease development. The seemingly lax study of the SuStaIn biotypes partially stems from the lack of large observational studies of the brain and behavior that would allow continued use and validations. The literature describes the frontal lobe, and it is shown to be a late maturing brain area and a very active area of synaptic changes, myelination changes, and anatomical changes; hence, making the lack of pattern to such a large portion of participants a potential sign of delayed neurodevelopmental characteristics which would putatively make it ideal and therefore our control group that does not have risk for executive and cognitive control dysfunction.

The Adolescent Brain Cognitive Development study aims to provide researchers with an unprecedented longitudinal dataset to determine novel insights about the trajectories related to the future development of mental health or substance use. In this work, we analyzed a biotype stratification using the SuStaIn model within the ABCD dataset and whether these show any association with behavior.

## Material and Methods

### Subjects

The selection of data is based on the availability of ABCD (Release 5.1). For our experiment, after an initial excluding criterion to remove subjects with neurological conditions, abnormal findings on brain scans, missing data, and siblings resulted in 8310 subjects. From these 8310 subjects, we selected 1368 subjects that have neuroimaging records on multiple time points, including baseline scans, 3rd-year scans, and 4th-year scans. This allows us to utilize delay discounting measurement on the tabulated data (nc_y_ddis.csv), which is only included in the 3rd year and 4-th year scans. The selection principle lies in a manageable balance subtype model complexity for SuStaIn and the possibility of potential longitudinal inquiries.

### Tabulated data

To focus on the frontal lobe development during adolescence, we consider the cortical thickness measurement on 20 frontal cortex ROIs out of 71 available ones on the tabulated data mri_y_smr_thk_dsk.csv. This file contains previously tabulated data of cortical thickness measurements in mm from the caudal anterior cingulate, caudal middle frontal, inferior temporal gyrus, isthmus cingulate, lateral orbitofrontal cortex, medial orbitofrontal cortex, parsopercularis, parsorbitalis, rostral middle frontal gyrus, and frontal pole. We also used the Delay Discounting results to determine associations between groups in the ABCD dataset. We used the indifference points, which represent the points where the small-immediate amount is deemed to have the same subjective value as a large, delayed reward. The indifference point measures how participants assess delayed rewards.

### SuStaIn

Our machine learning modeling using the Python version of SuStaIn^17^, pySuStaIn, to determine biotypes of neurodevelopmental trajectories. The pipeline can be summarized as follows:

1. Normalization of measurement by converting measurements into z-scores.
2. Estimate possible subtypes depending on staging inferences.
3. Verifying the optimal subtype model by cross-validation test.
4. Group the non-staging subjects as subtype-0 for a potential control group.

pySuStaIn runs on basic Python packages, including Numpy, Scipy, Scikit-learn, and Pathos. In our experiment, there were no objective control subjects, so we normalized the thickness data by the mean-thickness of all subjects included in the experiment. The cross-validation test can observe and suggest a balanced subtype model between model accuracy and complexity. Based on this suggested model selection, the non-staging subjects are grouped as subtype-0, i.e., these subjects do not contribute to any staging inference by SuStaIn.

### Statistics

Once the SuStaIn the subtype model selection, we carried out two statistical comparisons. These two comparisons use basic R packages and functions that enable ggplot() in R-studio. First, we calculated t-tests between subtype-0 and other subtypes to determine any behavioral difference between controls and biotypes. Then, to assess the association between frontal cortical thickness and related behavior, we performed correlations between the thickness of several interested frontal cortex ROIs and delay discounting measurements.

## Results

Once thickness measurements of the selected subjects were converted into the z-scores, we initialized the simulation to detect six subtypes. SuStaIn was able to cluster the subjects about four collective distributions within specific value ranges of Log-likelihood (Figures 1 and 2), indicating the potential reasonable number of subtypes could be four but not six because subtypes 5 and 6 do not distribute distinguishably from subtype 4 in terms of Log-likelihood values. To confirm this observation, we further carried out a 4-fold cross-validation test. This cross-validation verified that, by computing the Cross-Validation Information Criterion (CVIC) values, a balanced model selection for this dataset in between the model accuracy and model complexity should be four subtypes (Figures 3 and 4). The figures show that increasing the number of subtypes to more than four would not significantly improve model accuracy (a lower CVIC value and a higher Log-likelihood value). So, we continued our analysis with four subtypes (Figures 5 and 6). Eventually, those subjects non-staging subjects among subtypes 1 to 4 are categorized as subtype-0 and defined as control subjects (Table 1). In contrast, the staging subjects in subtypes 1 to 4 are classified as phenotype (Figure 7). Each biotype (B) identifies regions of the initial stages of neurodevelopment. The first biotype (B1) identifies the caudal anterior cingulate - isthmus cingulate - medial orbitofrontal as areas of interest for the neurodevelopment of this SuStaIn subtype. The second biotype (B2) identifies the lateral orbitofrontal cortex – parsorbitalis - medial orbitofrontal - frontal pole. The B3 identifies the caudal middle frontal - rostral middle frontal – parsopercularis. The B4 identifies the inferior temporal gyrus - isthmus cingulate. A significant portion (47%) of subjects cannot be associated with any of the four biotypes previously discussed, so they are identified as a last cohort that shows a lack of defined patterns of frontal lobe development, which we term a “control” subtype.

**Table 1.** Sustain identified cohorts.

| SuStaln biotype | Number of subjects |
| --- | --- |
| 0 (controls) | 660 |
| 1 | 215 |
| 2 | 205 |
| 3 | 158 |
| 4 | 130 |

**Figure 1.**
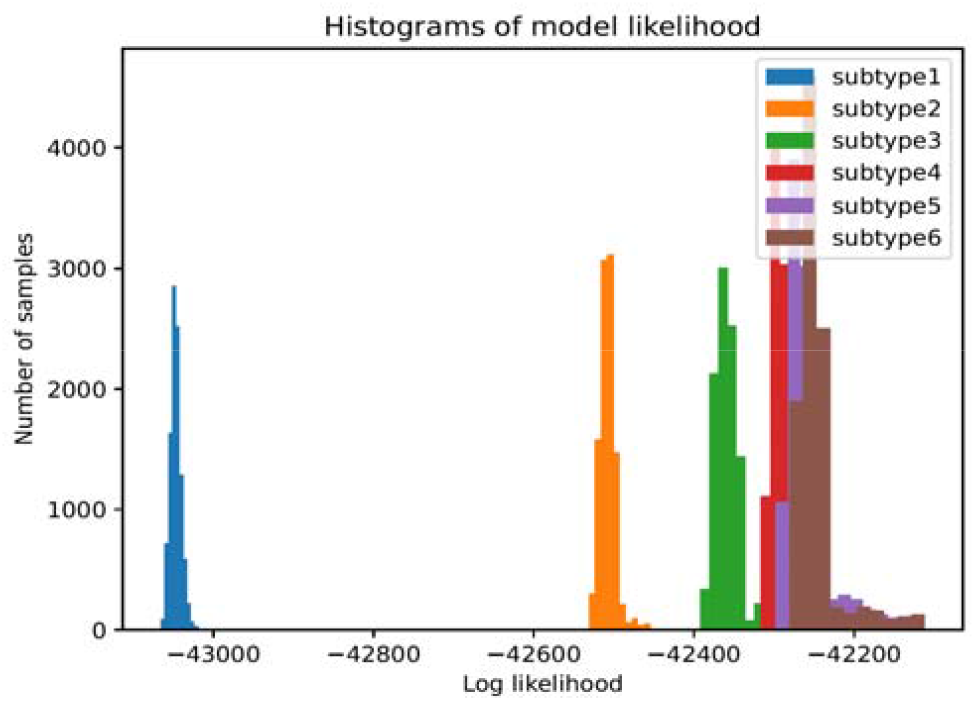
SuStaIn modeling likelihood of subtype identification for six subtypes. Significant overlap between subtypes four through six.

**Figure 2.**
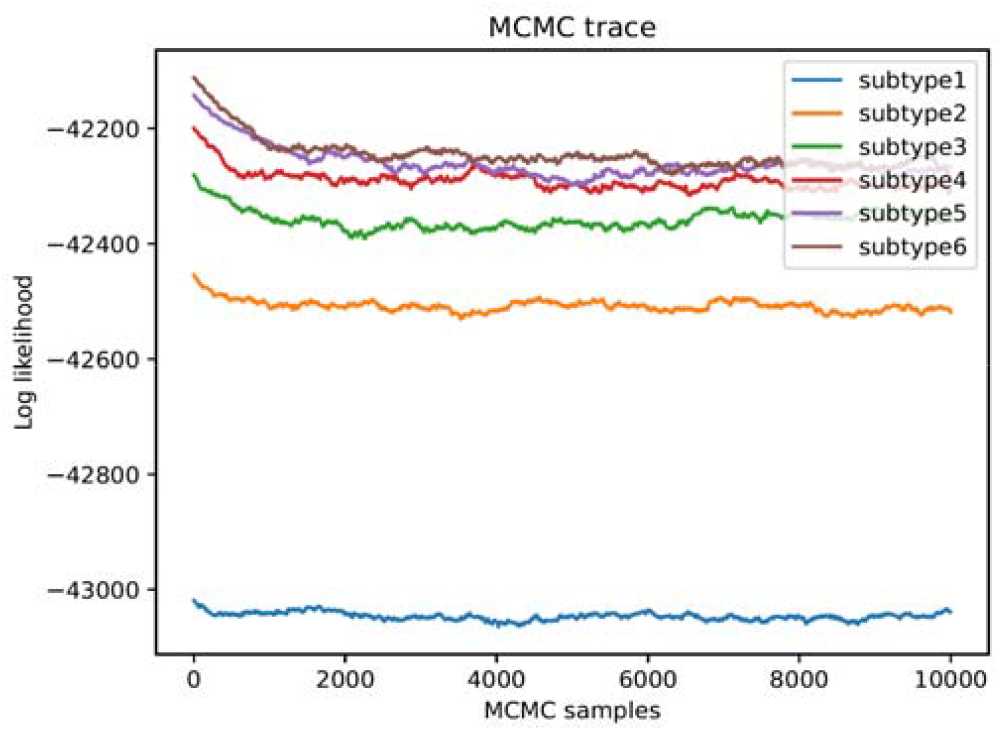
Markov-Chain Monte Carlo trace of the likelihood of SuStaIn modeling for six subtypes.

**Figure 3.**
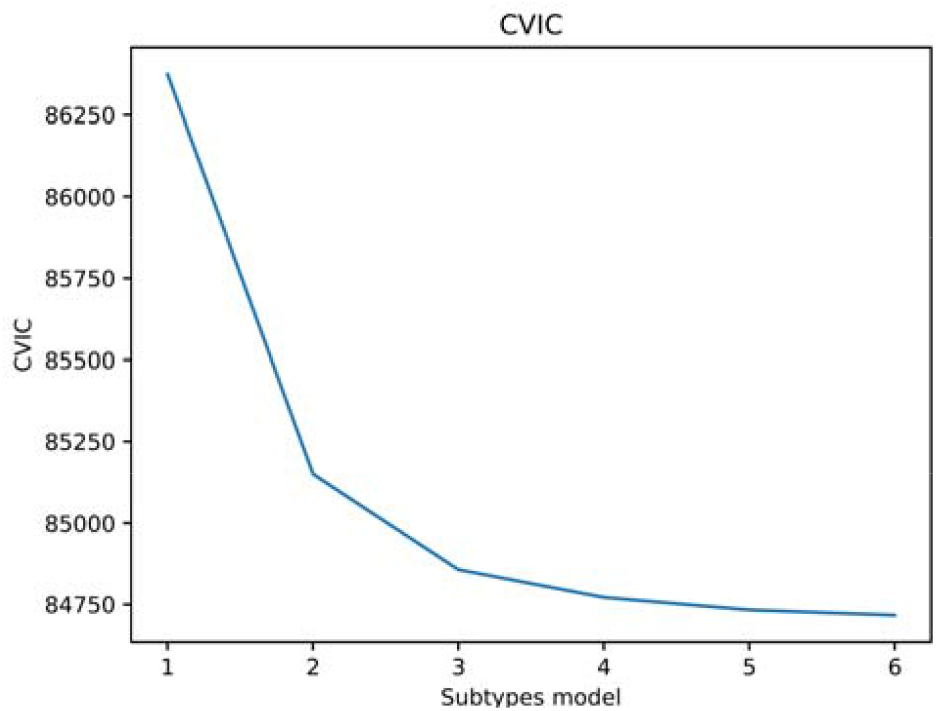
Cross validation curve for a SuStaIn simulation for six groups. Past the fourth subtype no significant subtypes are identified

**Figure 4.**
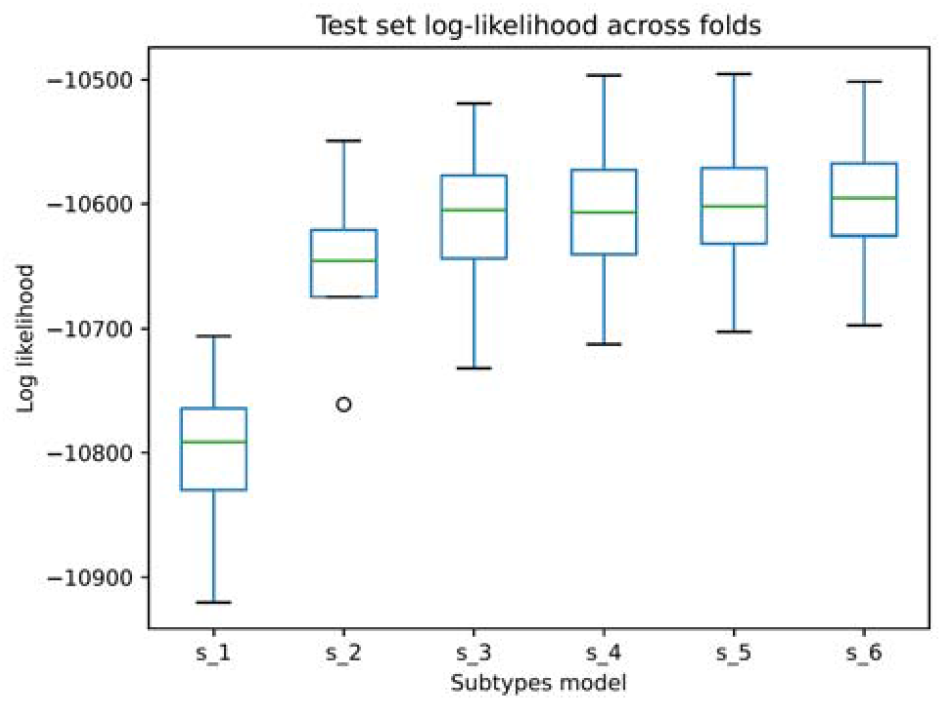
SuStaIn validation for six subtypes.

**Figure 5.**
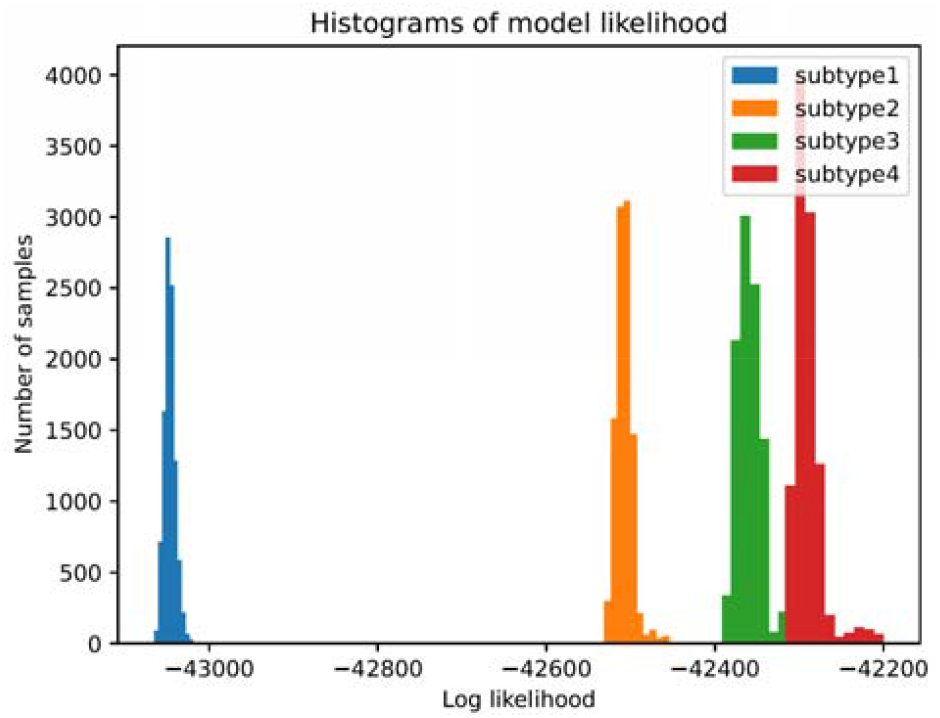
SuStaIn modeling likelihood of subtype identification for four subtypes show no overlap and succesful identification of four distinct biotypes.

**Figure 6.**
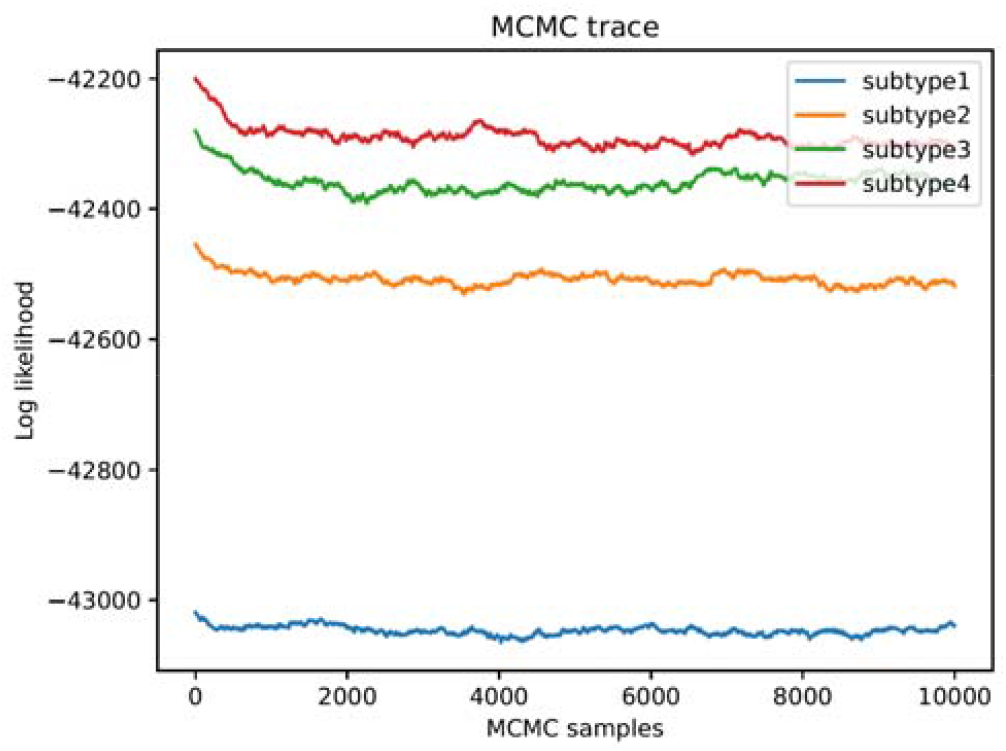
Markov-Chain Monte Carlo trace of the likelihood of SuStaIn modeling for four subtypes show no overlap and clearly defined distinct traces.

**Figure 7:**
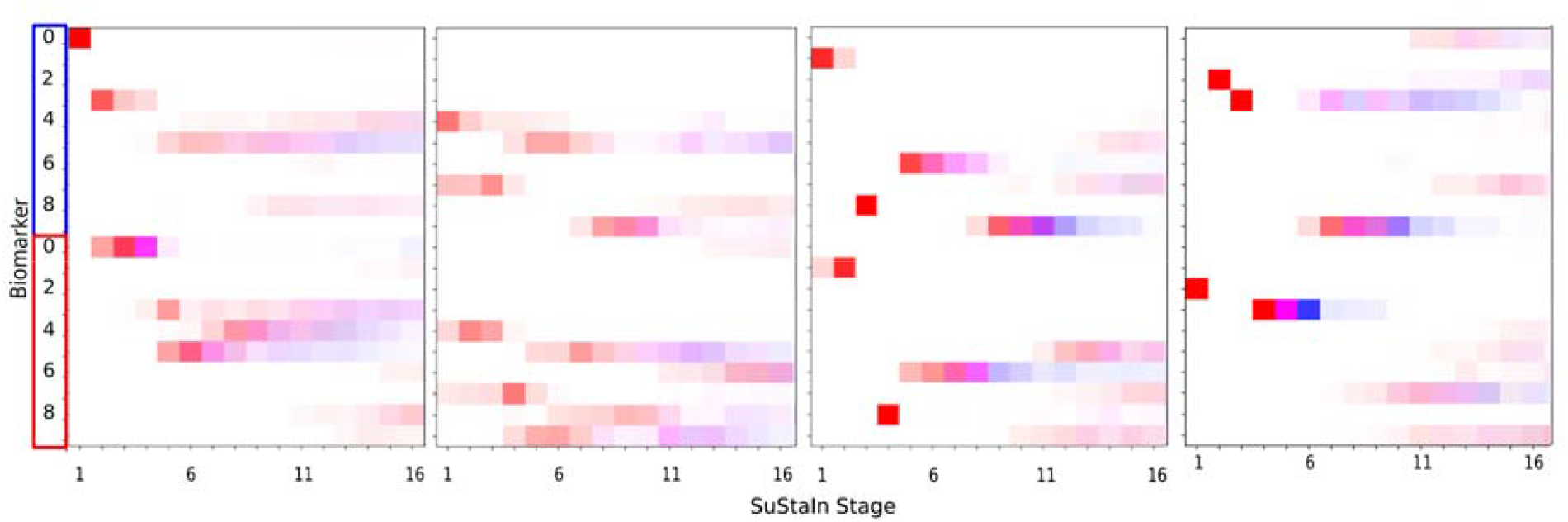
Subtypes and staging inferences for the resulting four distinct biotypes.

Based on the grouping of Table 1, the t-tests between the controls and each subtype, subtype-1 tends to show a trend of a reduction on the 3-month indifference point (p=0.067, Figure 8). Subtype 2 significantly decreases the 1-month indifference point (p=0.017, Figure 9). Subtype 3 did not display any reductions (Figure 10). Subtype 4 displays reductions in the 3- month indifference point (p=0.0092) and the 1-year indifference point (p=0.00418, Figure 11).

**Figure 8.**
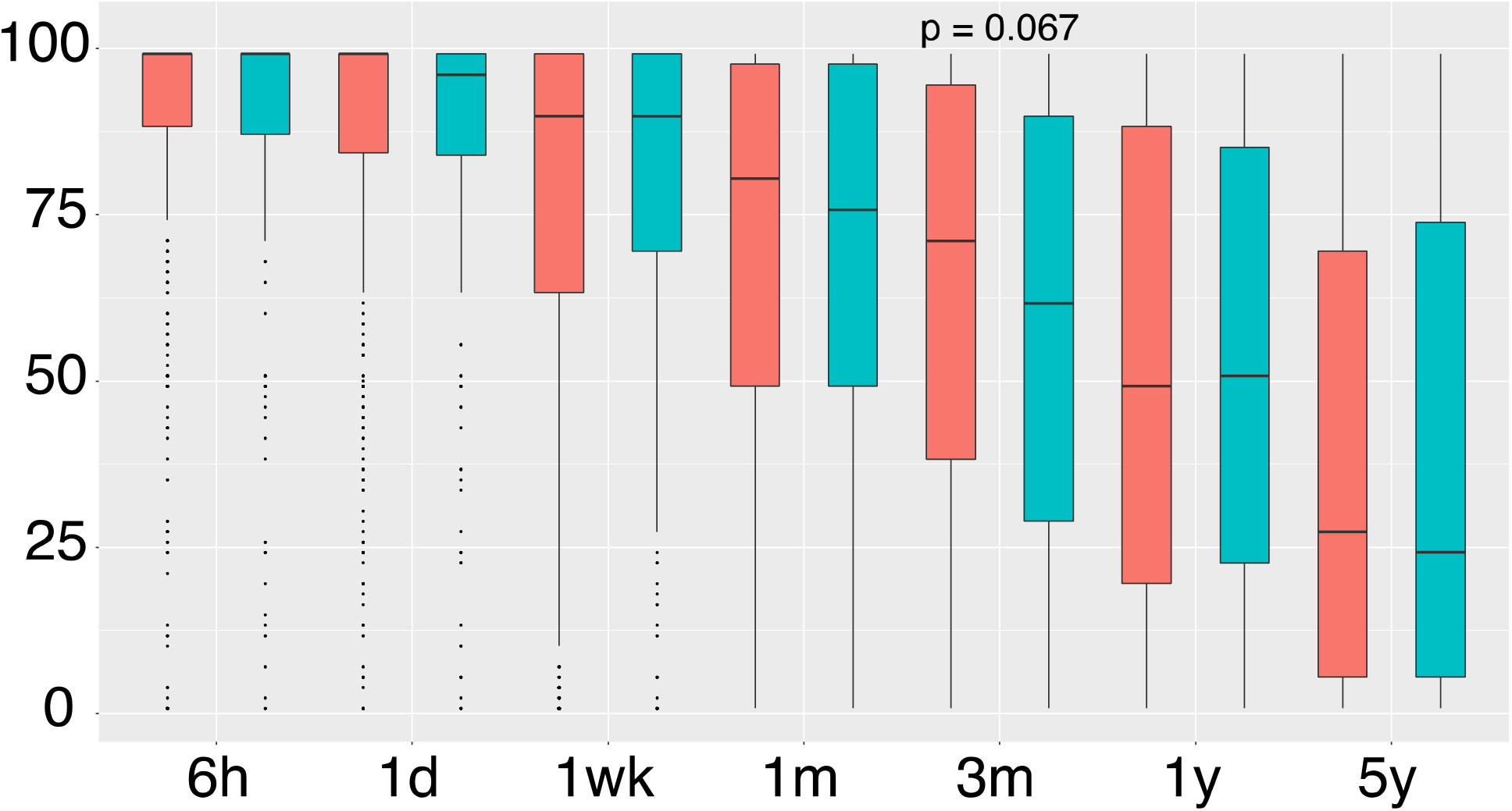
Delay discounting results for subtype 1 vs controls.

**Figure 9.**
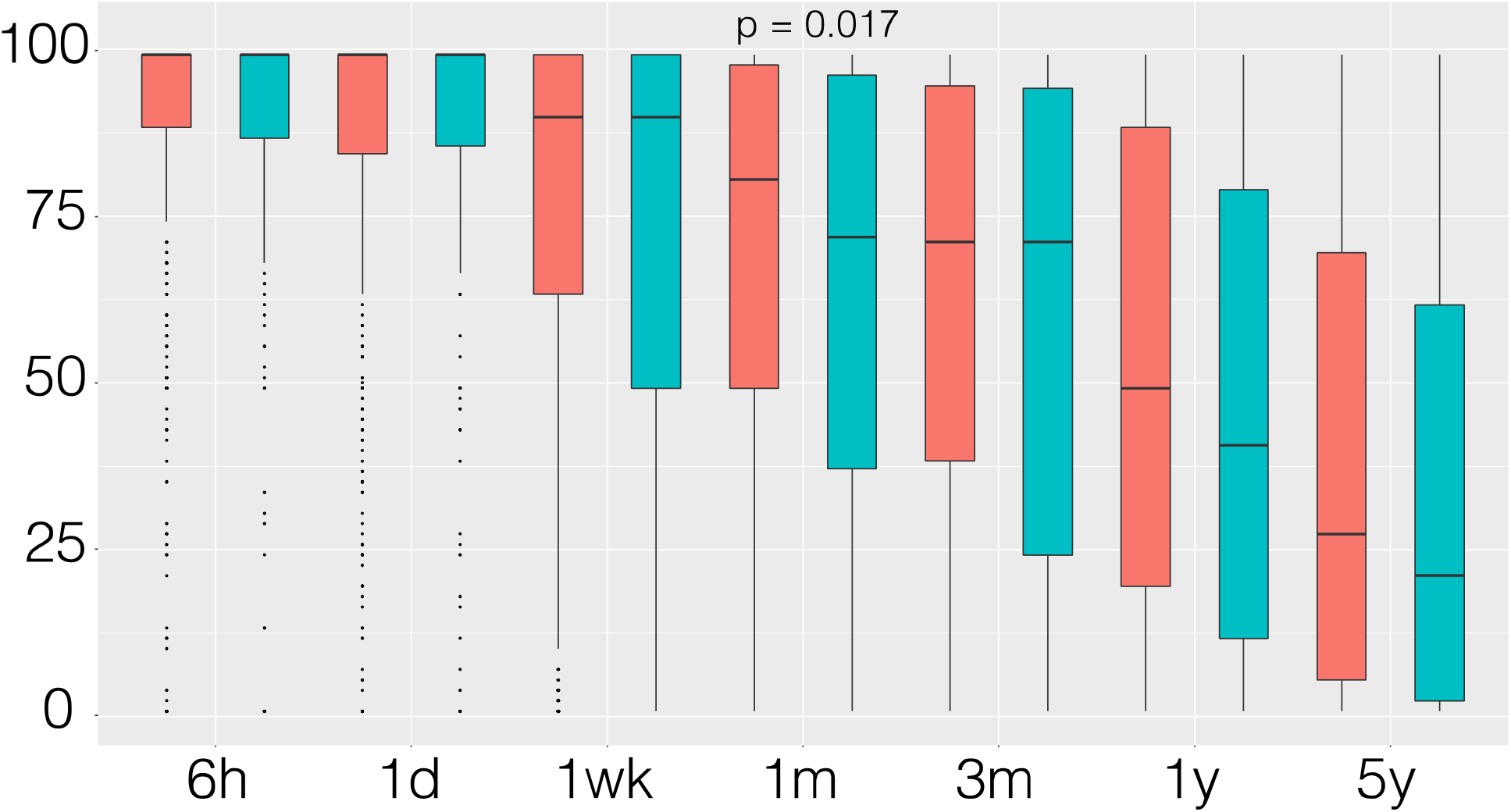
Delay discounting results for subtype 2 vs controls.

**Figure 10.**
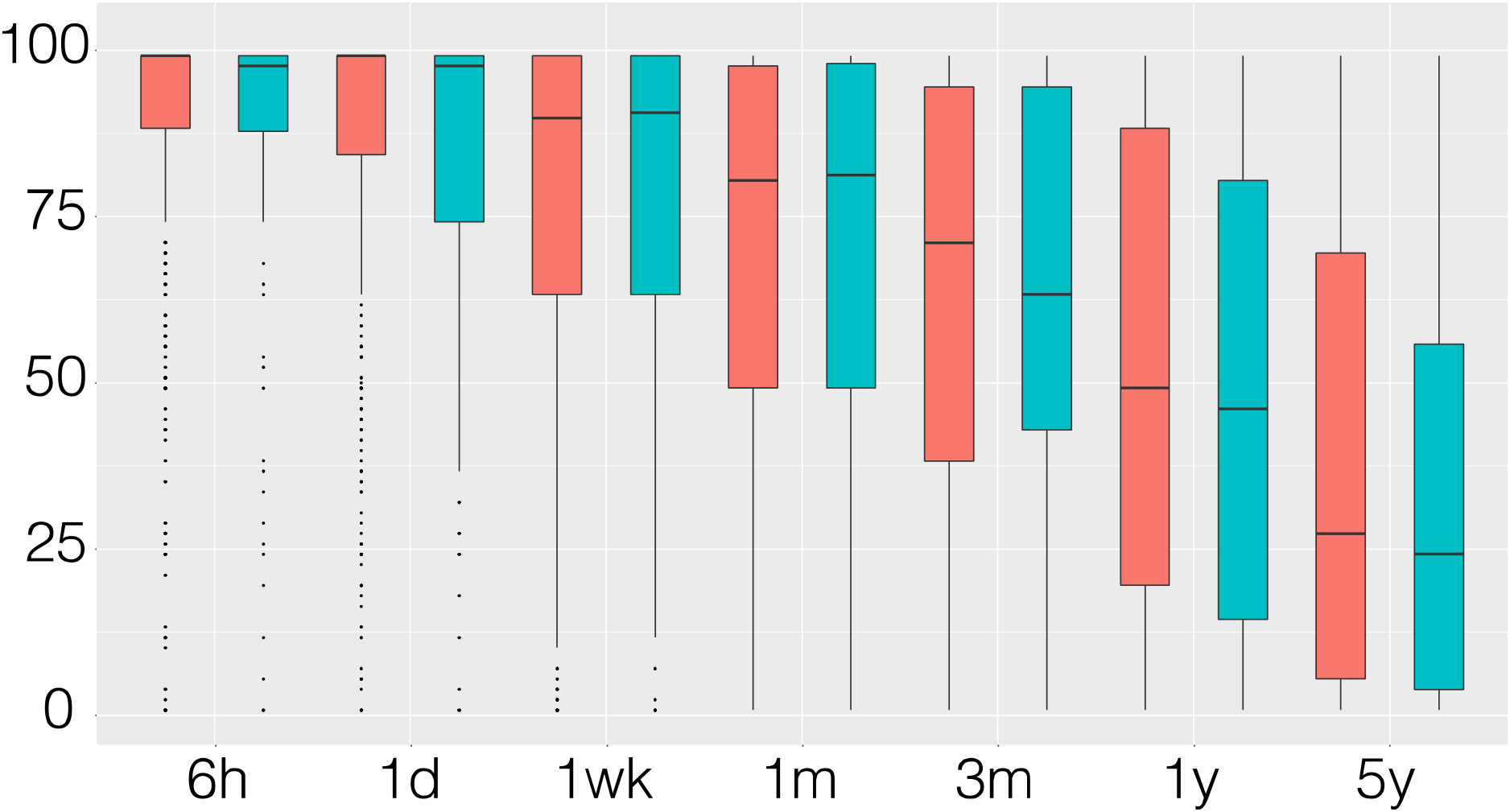
Delay discounting results for subtype 3 vs controls. No statistical differences found

**Figure 11.**
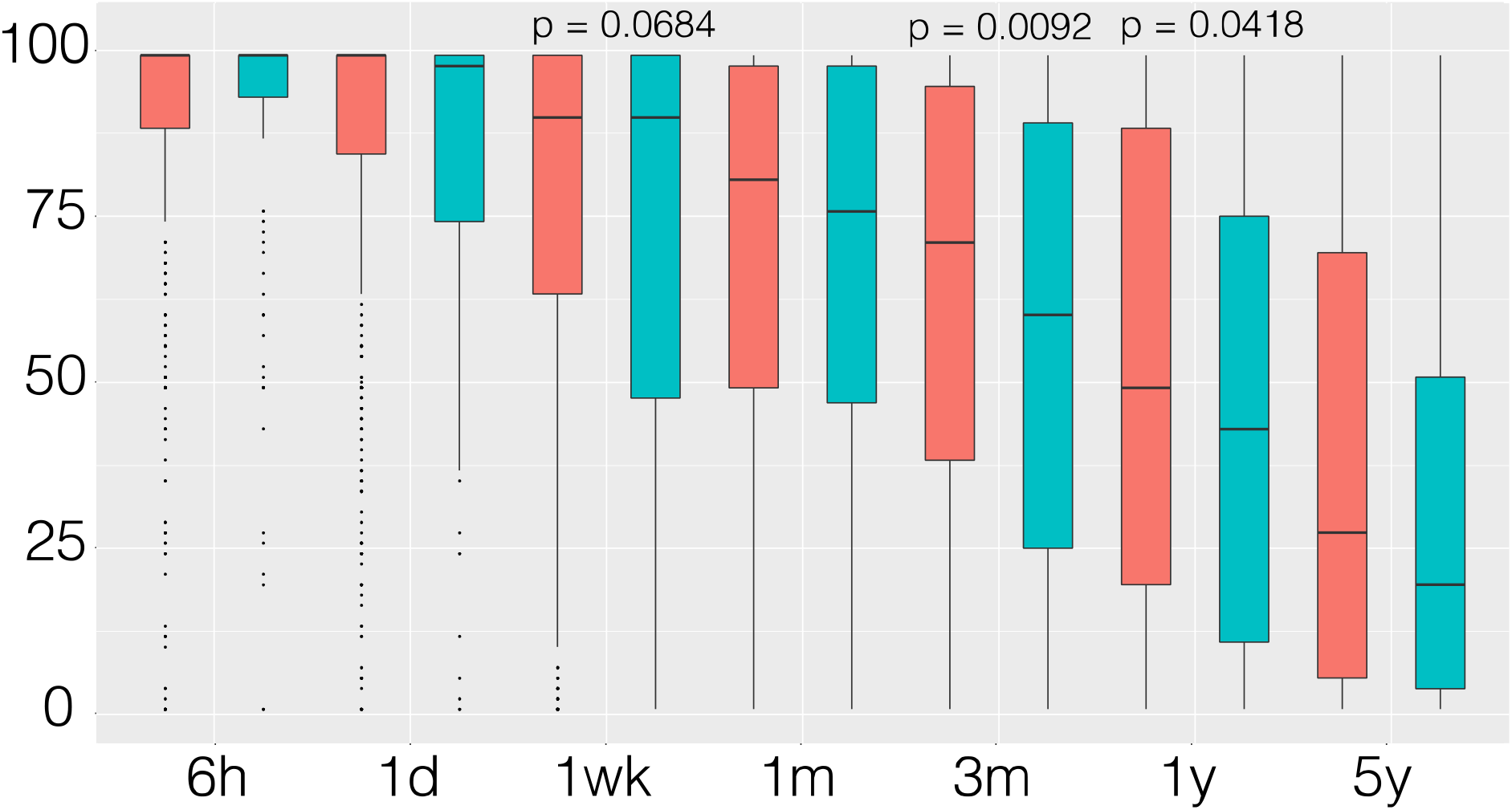
Delay discounting results for subtype 4 vs controls.

We then calculated correlations between each selected ROI. Only subtype 1 showed a significant correlation (R=-0.18, p=0.0095) between the thickness of left hemisphere caudal-anterior-cingulate (Figure 12) but nothing for the controls, which agrees with the tendency on the t-test on controls vs. subtype-1 (Figure 8). Such an association was not found for other subtypes (Figures 13-15).

**Figure 12.**
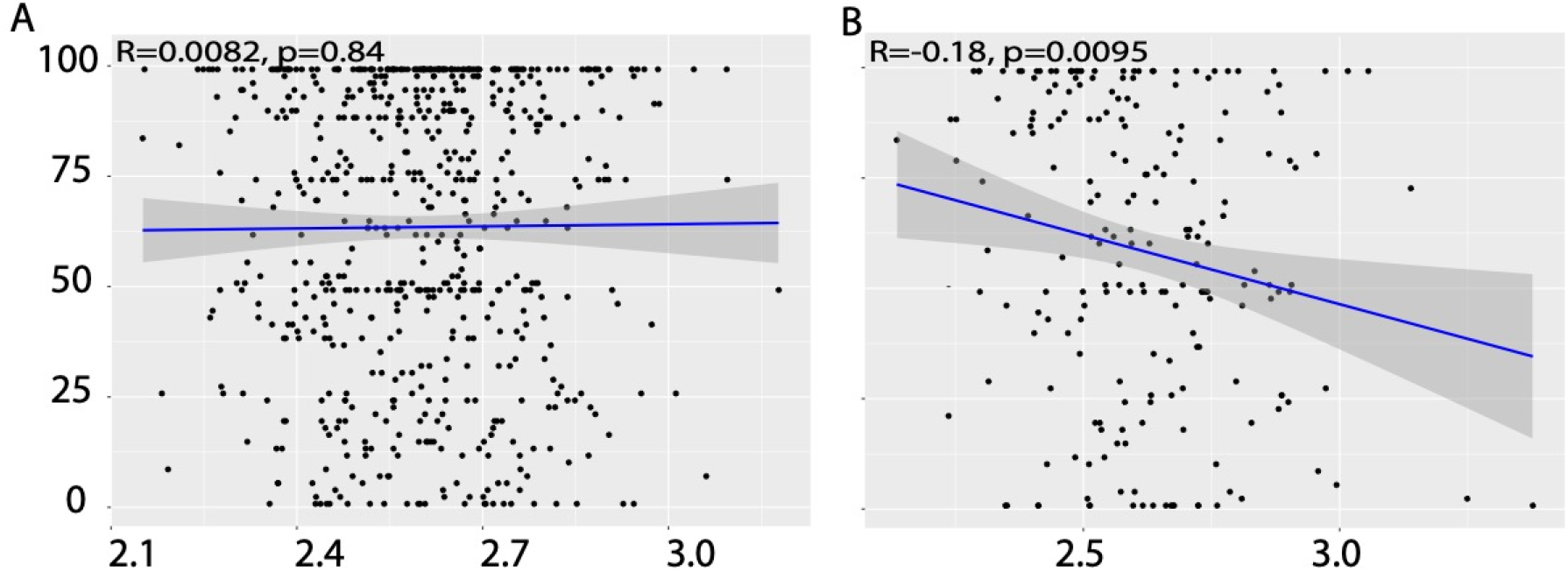
Correlation between the cortical thickness of the main feature of subtype 1 (See Fig 7) and delay discounting indifference point.

**Figure 13.**
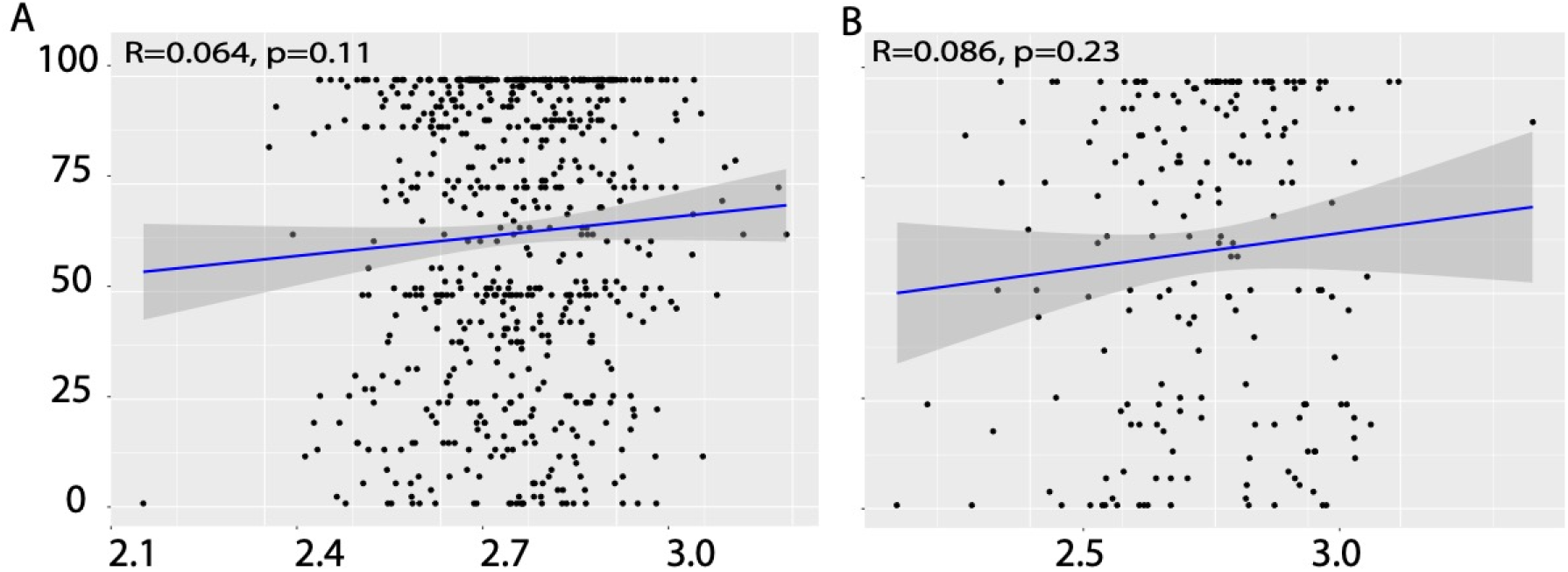
Correlation between the cortical thickness of the main feature of subtype 2 (See Fig 7) and delay discounting indifference point.

**Figure 14.**
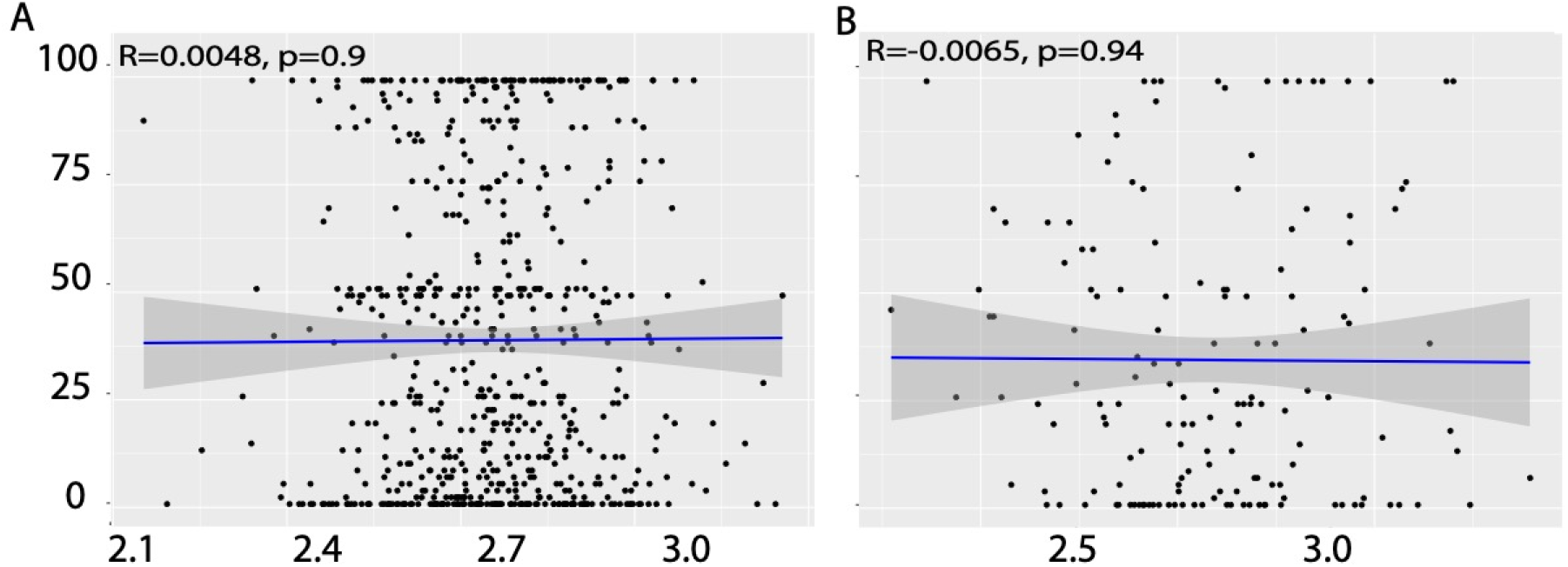
Correlation between the cortical thickness of the main feature of subtype 3 (See Fig 7) and delay discounting indifference point.

**Figure 15.**
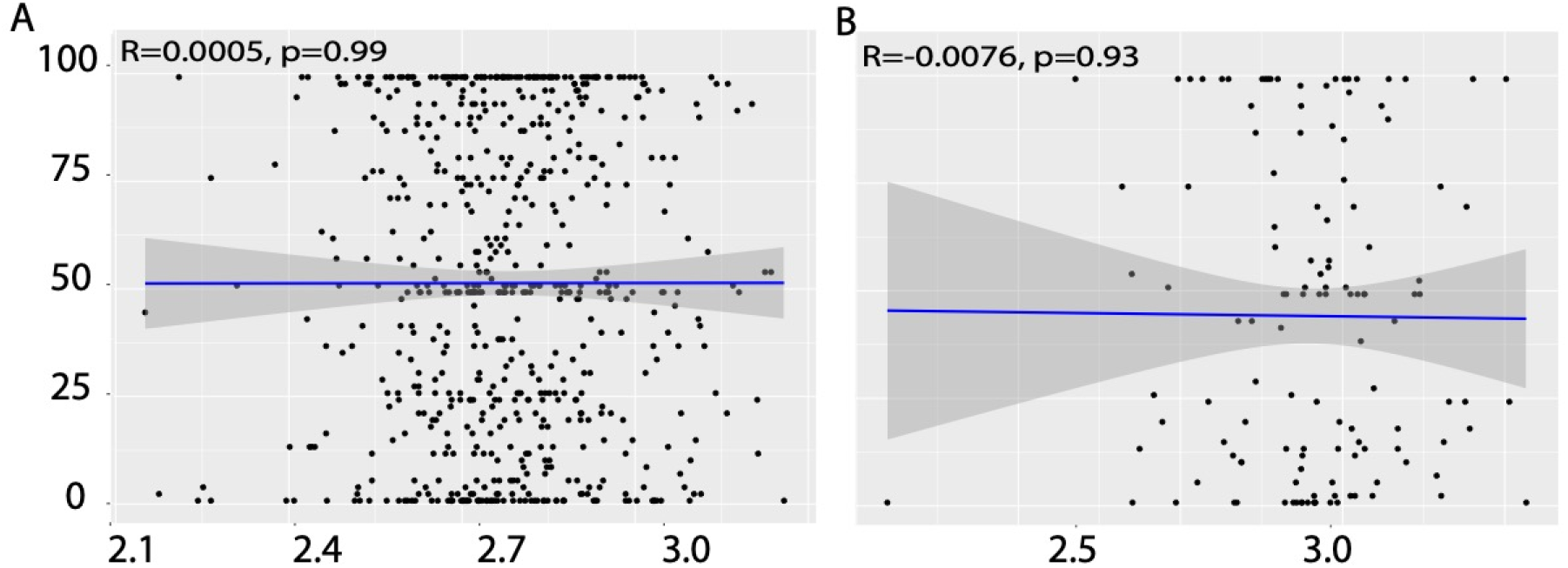
Correlation between the cortical thickness of the main feature of subtype 4 (See Fig 7) and delay discounting indifference point.

## Discussion

We successfully determined an initial biotype stratification of the ABCD dataset using SuStaIn modeling. We focused on cortical thickness because it is a recognized marker of pathology and proper development. Since thickness has been suggested as a proxy for worsening neurobiology, we continue with thickness as our biomarker. In the case of ABCD, a control cohort is not defined since the focus of the ABCD study is to find mental health trajectories and subsequent determination of risk and biomarkers. We maintain this principle and employ SuStaIn in a non-biased way by normalizing and converting z-score features by the mean and standard deviation of the features of all included participants, and we were able to identify four subtypes.

The literature describes the frontal lobe as a late maturing brain area and a very active area of synaptic changes, myelination changes, and anatomical changes; hence, making the lack of pattern to such a large portion of participants a potential sign of delayed neurodevelopmental characteristics which would putatively make it ideal and therefore our control group that does not have risk for executive and cognitive control dysfunction. The biotypes are accelerated patterns of neurodevelopment that show up at age nine and will serve as biomarkers of future executive dysfunction and, ultimately, drug use. The lack of it is a sign of adequate slow-paced frontal lobe development that will protect against executive dysfunction and, ultimately, drug use.

SuStaIn successfully identified a phenotypic subtype based on brain-derived frontal lobe features. We hypothesize that these subtypes are biomarkers of altered neurodevelopment that will be associated with domains of executive function or cognitive control. ^18^ The frontal lobe development is thought to extend up to the age of 25; however, this protracted development is not to be assumed that early during development (from age 9 to 19, ages that will be covered in the ABCD study) is mainly inactive. It is quite an active period that occurs as early as the second year of life.^19^ A detailed picture of the frontal lobe changes in anatomical volumetrics features (i.e., cortical thickness or regional volumes, myelin content, or gyrification characteristics) is not well defined, and most importantly, what early neurodevelopmental features relate to future characteristics in neurocognition, which we were able to confirm on an initial subset as an impulsive phenotype.

## Conclusion

Our preliminary analysis has identified biotypes in the ABCD dataset using SuStaIn. Initial results show that around half of the subjects do not show any apparent association with the biotype or stage, which we postulate to be a control cohort due to the heterogeneity of development in the protracted trajectories of the frontal lobe. Since slow-paced trajectories are associated with healthier behavioral development, we term these as controls. The four biotypes are associated with trajectories of specific brain region developmental features. Three of the biotypes show changes in delay discounting. Still, only one (B1) shows that cortical thickness of the caudal anterior cingulate (its first distinctive feature) correlates to delay discounting performance. In future work, we will fully characterize the anatomical features of these subtypes to longitudinally determine alterations in executive function and cognitive control and the association between pace and changes in several metrics of cognition.

## ABCD Acknowledgement

Data used in the preparation of this article were obtained from the Adolescent Brain Cognitive DevelopmentSM (ABCD) Study (https://abcdstudy.org), held in the NIMH Data Archive (NDA). This is a multisite, longitudinal study designed to recruit more than 10,000 children age 9-10 and follow them over 10 years into early adulthood. The ABCD Study® is supported by the National Institutes of Health and additional federal partners under award numbers U01DA041048, U01DA050989, U01DA051016, U01DA041022, U01DA051018, U01DA051037, U01DA050987, U01DA041174, U01DA041106, U01DA041117, U01DA041028, U01DA041134, U01DA050988, U01DA051039, U01DA041156, U01DA041025, U01DA041120, U01DA051038, U01DA041148, U01DA041093, U01DA041089, U24DA041123, U24DA041147. A full list of supporters is available at https://abcdstudy.org/federal-partners.html. A listing of participating sites and a complete listing of the study investigators can be found at https://abcdstudy.org/consortium_members/. ABCD consortium investigators designed and implemented the study and/or provided data but did not necessarily participate in the analysis or writing of this report. This manuscript reflects the views of the authors and may not reflect the opinions or views of the NIH or ABCD consortium investigators.

## Data availability

Data can be obtained through an NDA with NIMH. Instruction and relevant information can be found https://abcdstudy.org/scientists/data-sharing/

